# Faster sequence alignment through GPU-accelerated restriction of the seed-and-extend search space

**DOI:** 10.1101/007641

**Authors:** Richard Wilton, Tamas Budavari, Ben Langmead, Sarah Wheelan, Steven L. Salzberg, Alex Szalay

## Abstract

**Motivation:** In computing pairwise alignments of biological sequences, software implementations employ a variety of heuristics that decrease the computational effort involved in computing potential alignments. A key element in achieving high processing throughput is to identify and prioritize potential alignments where high-scoring mappings can be expected. These tasks involve list-processing operations that can be efficiently performed on GPU hardware.

**Results:** We implemented a read aligner called A21 that exploits GPU-based parallel sort and reduction techniques to restrict the number of locations where potential alignments may be found. When compared with other high-throughput aligners, this approach finds more high-scoring mappings without sacrificing speed or accuracy. A21 running on a single GPU is about 10 times faster than comparable CPU-based tools; it is also faster and more sensitive in comparison with other recent GPU-based aligners.

**Availability:** The A21 software is open source and available at https://github.com/RWilton/A21.

**Supplementary information:** Supplementary results are available at <<<TBD>>>

## 1 INTRODUCTION

The cost and throughput of DNA sequencing have improved rapidly in the past several years (Glenn, 2011), with recent advances reducing the cost of sequencing a single human genome at 30-fold coverage to around $1,000 (Hayden 2014). It is increasingly common for consortia, or even individual research groups, to generate sequencing datasets that include hundreds or thousands of human genomes. The first and usually the most time-consuming step in analyzing such datasets is read alignment. A read aligner will, for each sequencing read, attempt to determine its point of origin with respect to a reference genome. The continued dramatic growth in the size of sequencing datasets creates a crucial need for efficient and scalable read alignment software.

To address this need, a number of attempts have been made to develop read-alignment software that exploits the parallel processing capability of general-purpose graphics processing units, or GPUs. GPUs are video display devices whose hardware and system-software architecture support their use not only for graphics applications but also for general purpose computing. They are well suited to software implementations where computations on many thousands of data items can be carried out independently in parallel, and several high-throughput read aligners that use GPU acceleration have been developed in the past few years.

Experience has shown, however, that it is not easy to build useful GPU-based read alignment software. The salient problem is that the most biologically relevant sequence-alignment algorithm (Smith and Waterman, 1981; Gotoh, 1982) involves dynamic programming dependencies that are awkward to compute efficiently in parallel. For this reason, developers of read-alignment software have traditionally focused on optimized parallel implementations of the Smith-Waterman-Gotoh algorithm (Carriero and Gelernter, 1990; Manavski and Valle, 2008; Liu et al, 2013). Although the algorithm has been adapted to cooperative parallel-threaded GPU implementations (Khajeh-Saeed et al, 2010), the fastest GPU implementations of the algorithm have relied on task parallelism, where each thread of execution computes an entire pairwise alignment independently of all other parallel threads.

There is, however, another significant barrier to the implementation of high-throughput GPU-based alignment software. In a typical pairwise sequence alignment problem, a short query sequence (100 to 250bp) must be aligned with a comparatively long (10^9^ bp or longer) reference sequence. Since a brute-force search for all plausible alignments in this setting would be computationally intractable, read aligners typically construct a “search space” (a list of reference-sequence locations) within which potential alignments might be discovered. This aspect of the sequence alignment problem can account for a significant proportion of the computational effort involved in read alignment.

### 1.1 Seed and extend

The best-known algorithmic approach to exploring a reference-sequence search space is known as “seed and extend” (Lipman and Pearson, 1985).

**Figure 1.**
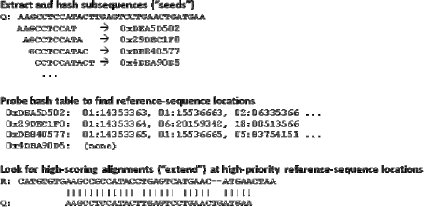
Seed-and-extend strategy for identifying potential alignments. Fixed-length subsequences (“seeds”) are extracted from the query sequence and hashed. Each hash value (e.g., “0xDEA5D502”) is used to probe a lookup table of reference locations (e.g., “01:14353363” for chromosome 1, offset 14353363) where the corresponding seed occurs. These seed locations are prioritized and full alignments between the query sequence and the reference sequence are explored in priority order.

An aligner that uses seed-and-extend relies on a precomputed index or lookup table to identify locations in the reference where a subsequence (“seed”) extracted from the query sequence matches the same-length subsequence in the reference. The aligner then performs a sequence-alignment computation at one or more of the reference-sequence locations it has obtained from the lookup table. In effect, the partial alignment implied by the seed match at each reference location is “extended” to arrive at a full pairwise alignment between the query sequence and the reference sequence.

### 1.2 Frequency distribution of seed locations

Most seed sequences occur rarely in the human reference genome, but a few seed sequences inevitably occur at hundreds or thousands of different locations in the reference sequence. This is not only because certain portions of the reference are internally repetitive (e.g., homopolymers or tandem repeats) but also because short sequences occasionally occur at two or more non-overlapping positions in the reference genome (e.g., because of transposon-induced duplication). This can be illustrated for the human reference genome by plotting the frequency with which 20mers (20bp subsequences) occur (Figure 2).

**Figure 2.**
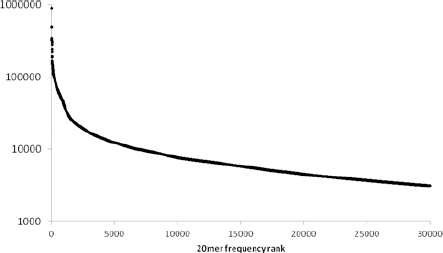
The number of different positions at which the 30,000 most frequently repeated 20mers occur in the human reference genome, ranked in descending order.

Although the mean frequency of human 20mers is only 10.7, high-frequency 20mers account for a disproportionate percentage of reference-sequence locations in a lookup table. For example, only 0.1% of the 20mers in the human reference genome appear in 200 or more different locations, but they account for about 10% of the 20mers in the reference sequence. In contrast, 71.7% of the 20mers in the human genome occur exactly once.

For a read aligner that implements a seed-and-extend strategy, this long-tailed distribution of seed frequencies is a computational obstacle for reads that contain one or more “high-frequency” seeds. To avoid searching for potential alignments at an inordinate number of locations in the reference sequence, an aligner must limit the number of locations at which it computes alignments.

### 1.3 Limiting the search space

Read aligners address this problem by using several heuristics, all of which limit the number of potential alignments computed:

- Limit the number of high-scoring mappings reported per read.
- Limit the number of seeds examined per read.
- Limit the number of reference locations examined per seed.

These heuristics trade throughput for sensitivity. An aligner spends less time computing potential alignments simply because it does not examine the entire search space (all reference locations for all seeds in each read). For the same reason, however, the aligner is less likely to identify all of the high-scoring mappings for each read. The A21 aligner implements two different heuristics to mitigate this problem.

#### 1.3.1. Reference-location sampling

In highly repetitive regions of the human genome, adjacent, overlapping 20mers in the reference sequence hash to a large number of locations in the reference sequence. A repetitive region is typically associated with long hash-table lists (“big buckets”) because the 20mers corresponding to those lists refer to numerous repetitive regions elsewhere in the reference. The A21 lookup tables are constructed by sampling adjacent “big bucket” hash-table lists in repetitive regions so that only one such list in 10 contains a reference-sequence location that lies within the region (Figure 3).

This sampling strategy decreases the size of large hash-table lists. The tradeoff is that a read that potentially aligns to a particular repetitive region must be seeded in up to 10 adjacent locations to guarantee that a reference location within the region will be found in a hash-table list for the read.

**Figure 3.**
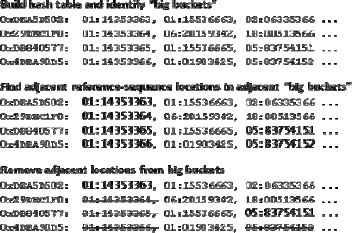
Adjacent reference-sequence locations are removed from the hash table when they are found in “big buckets” (hash-table lists whose cardinality exceeds a user-configurable threshold).

#### 1.3.2. Seed-coverage prioritization

At run time, A21 implements a heuristic that prioritizes alignments where a read contains two or more seeds that map to adjacent or nearby locations in the reference. This heuristic is reminiscent of the “spanning set method” used to compute alignments in the GSNAP aligner (Wu and Nacu, 2010), although A21 uses multiple neighboring seed hits only to identify high-priority locations for subsequent dynamic-programming alignment.

A21 uses an additional heuristic in the case of paired-end reads. The aligner prioritizes potential paired-end mappings where a reference location associated with at least one seed in one of the mates in the pair lies within a user-configurable distance and orientation of at least one seed in the other mate in the pair.

Notably, these heuristics are implemented using a series of sorting and reduction operations on an aggregated list of seed locations. In a CPU-based implementation, the amount of computation required for these operations would be impractical with a reference genome the size of the human genome. In a GPU-based implementation, however, these list-based operations can be performed efficiently with a combination of cooperative parallel threading (sort, stream compaction) and task parallelism (computing seed coverage, filtering using paired-end criteria). For example, an NVidia GTX480 GPU can sort over 300 million 64-bit integer values per second.

The A21 aligner was designed to evaluate the performance of these “GPU-friendly” heuristics. In effect, A21 implements a pipeline in which the following operations are performed on GPU hardware for each read:

- Define the “search space” for the read; that is, compute the set of reference locations that correspond to the seeds in the read.
- Adjust the reference locations so that they correspond to the location of the seed within the read.
- Sort the list of reference locations.
- For paired-end reads, identify pairs of reference locations that lie within a predefined distance and orientation of each other.
- Coalesce adjacent seed locations so that they are covered by a minimum number of alignment computations.
- Compute alignments to identify and record high scoring mappings.

## 2 METHODS

The A21 aligner is written in C++ and compiled for both Windows (with Microsoft Visual C++) and Linux (with the Gnu C++ compiler). The implementation runs on a user-configurable number of concurrent CPU threads and on one or more NVidia GPUs. The implementation pipeline uses about 30 different CUDA kernels written in C++ (nongapped and gapped alignment computation, application-specific list processing) and about 100 calls to various CUDA Thrust APIs (sort, reductions, set difference, string compaction).

The development and test computers were each configured with dual 6-core Intel Xeon X5670 CPUs running at 2.93GHz, so a total of 24 logical threads were available to applications. There was 144GB of system RAM, of which about 96GB was available to applications. Each computer was also configured with three NVidia Tesla series GPUs (Kepler K20c), each of which supports 5GB of on-device “global” memory and 26624 parallel threads. The internal expansion bus in each machine was PCIe v2.

Throughput (as query sequences aligned per second) was measured only when no other user applications were using the machines so that all CPU, memory, and I/O resources were available. For experiments with simulated data, we used Mason (Holtgrewe, 2010) to generate 1 million 100bp paired-end reads. For experiments with Illumina data, we used 100bp paired-end Illumina Genome Analyzer data from the YanHuang genome (Li et al, 2009).

### 2.1 Software implementation

The A21 implementation is a pipeline in which batches of reads are processed by a sequence of discrete software modules, each of which operates on a separate CPU thread that is allocated for the lifetime of the module and then discarded. When multiple GPUs are used, each GPU is associated with its own CPU thread. Modules are designed to execute concurrently on CPU threads and on the GPU. Data common to multiple CPU threads is shared; data common to a sequence of GPU operations resides in GPU device memory without being transferred to or from CPU memory.

**Figure 4.**
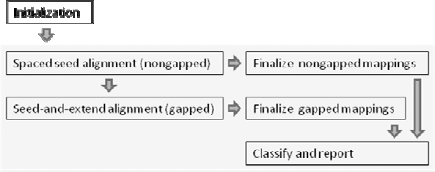
The A21 pipeline implementation consists of one-time-only initialization (memory allocation, loading of lookup tables and reference data) followed by iterative batched processing of reads (query sequences). Within each batch, nongapped alignments are discovered using GPU-based spaced seed alignment. Gapped alignments, using GPU-based seed-and-extend alignment, are computed only for reads for which a satisfactory number of nongapped alignments are not found. All mappings are finalized (scored and mapped), classified, and reported in multiple concurrent CPU threads.

The execution of the A21 pipeline consists of iterative processing of batches of reads (query sequences), where the number of reads in a batch is constrained by the amount of available GPU memory. Within each batch iteration, the GPUs are used for list processing and for the computation of alignments, while CPU threads are used concurrently for scoring, classification, and formatting of alignment results as well as for input and output. GPU code executes concurrently with CPU code wherever possible. For example, the classification, reporting, and final output of the alignment results for a batch overlaps with the beginning of processing of the subsequent batch.

### 2.2 Nongapped alignment

The nongapped aligner uses periodic spaced seeds to identify potential mappings (Chen et al., 2009). The seed value used in A21 covers 84 adjacent positions with 30 “care” positions. It is fully sensitive to nongapped alignments containing up to two mismatches when seven overlapping seed comparisons are used. Both the seed value and the query sequences are encoded in 64-bit packed binary values to facilitate bitwise operations between the seed value and the binary representation of each query sequence.

For each of the first seven positions in each query sequence, the result of the bitwise AND between the seed value and the query sequence is packed into a 30-bit value that is used to probe a lookup table of potential alignment locations in the reference sequence. For each such location, mappings between the query sequence and the reference sequence are identified by bitwise comparison of the entire query sequence with the corresponding reference.

Nongapped mappings with mismatches near one or both ends are examined for potential soft clipping. A21 soft-clips a nongapped mapping whenever its alignment score is higher than it would be without soft clipping. The nongapped aligner assigns a numerical score to each mapping by applying the user-specified parameters for Smith-Waterman-Gotoh affine-gap alignment.

### 2.3 Gapped alignment

A21 performs gapped alignment only on reads for which it does not find a sufficient number of nongapped alignments. The minimum number of satisfactory nongapped alignments required for a read to be excluded from further processing is a user-configurable parameter.

The gapped aligner is a straightforward implementation of the seed-and-extend strategy. To facilitate parallel computation, multiple seed locations are examined concurrently within each read. Groups of seed locations are selected iteratively. The first group of seeds is chosen so as to cover the entire read without overlapping seeds; subsequent groups are selected so as to overlap the seed positions examined in previous groups.

In each iteration, the seed subsequences are extracted from the read and hashed to 30 bits. The 30-bit hash values are used to probe a hash table of reference-sequence locations. The reference locations are prioritized and Smith-Waterman-Gotoh local alignment is computed at the highest-priority locations. Reads for which a user-configured number of satisfactory mappings have been found are excluded from subsequent iterations.

Each iteration examines seed locations that straddle the locations that were processed in previous iterations; seeds are chosen at locations that are halfway between those examined in all previous iterations. (This is similar to the behavior of Bowtie 2’s -R option.) In this way the cumulative number of seeds examined doubles with each iteration, but the actual number of reference locations considered remains relatively stable. Because A21 uses fixed-length 20bp seeds (20mers), six “seed iterations” are required to examine every seed location in the query sequence.

### 2.4 Restricting the seed-and-extend search space

To facilitate GPU-based list operations, the A21 implementation encodes reference locations as 64-bit bitmapped values that can be represented in one-dimensional arrays. These arrays are maintained exclusively in GPU device memory where multiple CUDA kernels can access them. CUDA kernels are used to reorganize and triage reference-location lists:

- Prioritize reference locations that lie within paired-end distance and orientation constraints.
- Prioritize reference locations where overlapping and adjacent seeds cover the largest number of adjacent positions in the reference sequence.
- Exclude reference locations that have been examined in previous seed iterations.
- Identify reference locations for which acceptable mappings exist and for which criteria for paired-end mapping are met.

### 2.5 Specific concerns for GPU implementation

Available memory and computational resources on GPU devices constrain the implementation of the A21 pipeline. Although the compiled code is not “tuned” to a particular GPU device, the source-code implementation follows programming practices that experience has shown lead to higher performance: judicious use of GPU memory and use of data-parallel algorithms and implementation methods.

#### 2.5.1. Memory size

The limited amount of on-device GPU memory constrains the amount of data that can be processed at any given time on a GPU. Because GPU memory requirements vary as data moves through the implementation pipeline, it is impossible to provide for full usage of available GPU memory at every processing step.

The approach taken in A21 is to let the user specify a batch size that indicates the maximum number of reads that can be processed concurrently. In computations where available GPU memory is exceeded (for example, in performing gapped local alignment), A21 breaks the batch into smaller sub-batches and processes the sub-batches iteratively.

#### 2.5.2. Memory layout

The A21 implementation pays particular attention to the layout of data in GPU memory. Memory reads and writes are “coalesced” so that data elements accessed by adjacent groups of GPU threads are laid out in adjacent locations in memory. A21 therefore uses one-dimensional arrays of data to store the data elements accessed by multiple GPU threads. Although this style of in-memory data storage leads to somewhat opaque-looking code, the improvement in the speed of GPU code is noticeable (sometimes by a factor of two or more).

#### 2.5.3. Minimal data transfers between CPU and GPU memory

Although data can theoretically move between CPU and GPU memory at speeds determined by the PCIe bus, experience has shown that application throughput is decreased when large amounts of data are moved to and from the GPU. For this reason, A21 maintains as much data as possible in GPU memory. Data is transferred to the CPU only when all GPU-based processing is complete.

#### 2.5.4. Divergent flow of control in parallel threads

Divergent flow of control in adjacent GPU threads can result in slower code execution. Branching logic is therefore kept to a minimum in GPU code in A21. Although this problem was encountered in previous GPU sequence-aligner implementations (Schatz et al, 2007), it is empirically less important in the A21 implementation than the effect of optimized GPU memory access.

### 2.6 Analysis of alignment results

We used the human reference genome release 37 (Genome Reference Consortium, 2014) for throughput and sensitivity experiments. We compared A21 results with the output generated by two widely-used CPU-based read aligners and two recent GPU-based aligners (software versions listed in Supplementary Table T1):

- Bowtie 2 (Langmead et al, 2013) (CPU)
- BWA-MEM (Li, 2013) (CPU)
- SOAP3-DP (Luo et al, 2013) (GPU)
- NVBIO (NVidia, 2014) (GPU)

We parsed the SAM-formatted output (SAM/BAM Format Specification Working Group, 2013) from each aligner and aggregated the results reported by each aligner for each read. We examined the POS (position), TLEN (paired-end fragment length), and AS (alignment score) fields to ensure the consistency of the set of mappings reported by each aligner. For SOAP3-DP, which does not report alignment scores, we derived scores from the mapping information reported in the CIGAR and MD fields. We used the following scoring parameters: match=+2; mismatch=-6; gap open=-5; gap space=-3, with a threshold alignment score of 100 (for 100bp reads) or 400 (for 250bp reads).

We used simulated (Mason) reads to evaluate sensitivity for both paired-end and unpaired reads. For each aligner, we used high “effort” parameters so as to maximally favor sensitivity over throughput. For each read mapped by each aligner, we compared the POS and CIGAR information reported by the aligner with the POS and CIGAR generated by Mason. We assumed that a read was correctly mapped when, after accounting for soft clipping, one or both of its ends mapped within 3bp of the mapping generated by Mason. (Supplementary Table T2 explains our choice of a 3bp threshold.) To illustrate sensitivity and specificity, we plotted the cumulative number of correctly-mapped and incorrectly-mapped reads reported by each aligner, stratified by the MAPQ score (Li et al, 2008) for each read.

We used the YanHuang data to measure throughput using both paired-end and unpaired reads. For this analysis, we recorded throughput across a range of “effort” parameters chosen so as to trade speed for sensitivity. We defined “sensitivity” as the percentage of reads reported as mapped by each aligner with alignment score (and, for paired-end reads, TLEN) within configured limits.

Prior to computing alignments, all of the GPU-aware aligners spend a brief period of execution time initializing static data structures in GPU device memory. We excluded this startup time from throughput calculations for these aligners.

## 3 RESULTS

Each of these read aligners is able to map tens of millions of reads to the human genome in an acceptably short period of time. All of the aligners, including A21, were capable of mapping reads with high accuracy. In terms of throughput, however, A21 demonstrated higher throughput than the other aligners across a wide range of sensitivity settings.

### 3.1 Evaluation on simulated data

With simulated Illumina read data, A21mapped paired-end reads to their correct origin in the reference genome with sensitivity and specificity comparable to all four of the aligners to which we compared it (Figure 5 and Supplementary Figures S1-S8). Although each aligner uses a slightly different computational model for MAPQ, all of the aligners maintain a very high ratio of correct to incorrect mappings until mappings with relatively low MAPQ scores are considered.

**Figure 5.**
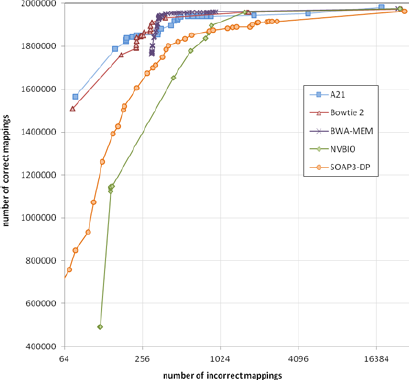
Total correctly mapped versus incorrectly mapped reads, plotted for decreasing MAPQ, for 1 million simulated 100bp paired-end Illumina reads (2 million total reads). Results for unpaired reads and for 250bp reads are very similar (Supplementary Figures S1-S8).

### 3.2 Evaluation on real data

We used experimental data from the YanHuang human genome project to evaluate speed (Figure 6 and Supplementary Figure S9).

**Figure 6.**
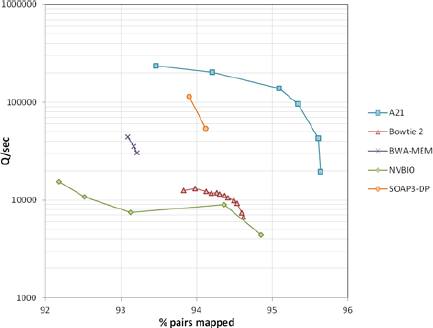
Speed (measured as the number of 100bp query sequences processed per second) plotted versus sensitivity (expressed as the overall percentage of mapped pairs). Data for 10 million 100bp paired-end reads from the YanHuang genome. Workstation hardware: 12 CPU cores (24 threads of execution), one NVidia K20c GPU. Results for unpaired reads are similar (Supplementary Figure S9).

Throughput decreases with increasing sensitivity for all of the aligners, with a more rapid decrease near each aligner’s maximum sensitivity. This is apparent even with BWA-MEM and SOAP3-DP, although we were unable to “tune” these aligners across as wide a range of sensitivity settings as the others.

A21’s speed on a single GPU is generally about 10 times that of the CPU-based aligners to which we compared it. When compared with GPU-based aligners, A21 is about 10 times faster than NVBIO and about twice as fast as SOAP3-DP. When executed concurrently on multiple GPUs in a single machine, A21’s throughput increases in proportion to the number of GPUs (Figure 7). At lower sensitivity settings, overall throughput is limited by PCIe bus bandwidth. Scaling improves at higher sensitivity settings, where throughput is limited by the number of dynamic-programming computations carried out on the GPUs.

**Figure 7.**
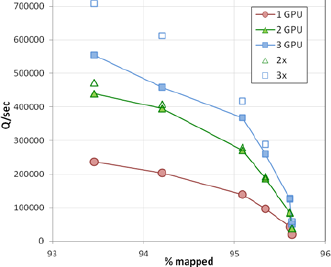
Throughput on one, two, and three GPUs (NVidia K20c) in a single computer for the data shown in Figure 6. For comparison, 2x and 3x multiples of single-GPU throughput are also plotted.

## 4 DISCUSSION

Apart from its potential for high throughput, the A21 implementation demonstrates that an increase in throughput can be achieved without losing sensitivity. Furthermore, by sacrificing throughput, A21 can be “pushed” to a comparatively high level of sensitivity.

## 4.1 Performance characteristics

It is clear from the shape of its speed-versus-sensitivity curve that A21 achieves increased sensitivity by exploring a proportionally larger search space per successful mapping. A21’s search-space heuristics cause it to find high-scoring mappings (that is, perfect or near-perfect alignments) rapidly within a relatively small search space. For reads that do not map with high alignment scores, however, A21 must explore more seed locations and compute more dynamic programming problems before it can report a satisfactory mapping. For example, in the experiment shown in Figure 6, A21 computed about 8 times as many dynamic-programming alignments at the high end of its sensitivity range as it did at the low end of the range.

A21 explores a significantly larger search space for reads that it cannot align with a comparatively small number of mismatches or gaps. This mitigates the effect of the heuristics that filter the list of potential mapping locations on the reference sequence. In particular, gapped mappings that might be missed in an early seed iteration (when seeds are spaced widely) are detected in later seed iterations (when seeds are spaced more closely). The nature of these heuristics, however, implies that the additional mappings that A21 finds when it is configured for higher sensitivity are generally lower-scoring mappings.

The effect of A21’s heuristics on the computation of MAPQ (mapping quality) for a read is difficult to determine. In some cases, A21 assigns a lower MAPQ (higher probability that the read is incorrectly mapped) simply because it computes alignments in parallel for the read and therefore tends to find more alternative mappings than would a non-parallelized implementation. On the other hand, by excluding many potential reference locations (and thus potential alternative mappings) from its search space, A21 might incorrectly assign a high MAPQ to a read. In any event, we do not observe any notable difference overall in A21’s MAPQ scoring when compared with other aligners.

## 4.2 GPU-accelerated sequence alignment

Unlike the Smith-Waterman-Gotoh alignment algorithm, parallel list-management algorithms — in particular, variations of radix sort and of prefix (scan) operations — are amenable to cooperative parallel-threaded implementation. Much of the intermediate processing in the sequence-alignment pipeline is thus well suited to GPU-based implementation. We exploited this characteristic of the read-alignment process in designing and developing the A21 software.

There are two features in the design and implementation of A21 that distinguish it from other GPU-based read aligners. First, we adopted a software-design approach that used the GPU as an “accelerator” in a task-parallel pipeline. CPU threads execute concurrently with GPU threads on independent data wherever possible, with synchronization points only where the GPU has completed processing a set of data. In practice, this means that overall throughput is GPU-bound and thus insensitive to variations in the time spent executing CPU threads (including post-processing alignments, reading and writing data files, and recording performance data).

Second, we used GPUs for the kinds of computations for which they are well suited (stream compaction, radix sort, parallel prefix reduction, set difference). This is a reasonable approach not only because it uses GPU hardware in a natural way, but also because we were able to leverage the NVidia Thrust library, one of several freely-available, well-optimized library implementations of basic parallel operations on GPU hardware. This approach led to an emphasis on data structures that can be represented in one-dimensional arrays of integers as well as list manipulations that involve simple, data-independent numerical operations.

There is a rule of thumb among GPU programmers that states that the cost and effort of developing software for GPUs is justified only when an order-of-magnitude improvement can be obtained relative to a comparable non-GPU implementation. Our experiments with A21 show that it meets this informal performance criterion.

We nevertheless recognize that direct comparisons in speed between CPU-based and GPU-based software implementations are fraught with difficulties (Anderson et al, 2011). We attempted to choose comparison hardware that was reasonably similar in terms of cost and availability. As more capable CPU and GPU hardware becomes available, we expect A21, like all of the aligners we evaluated, to deliver higher throughput.

We also foresee further optimization of A21’s implementation. For example, there are newer, faster, GPU function libraries that might be used to replace calls to the Thrust APIs. Also, we have not experimented with low-level optimization of our Smith-Waterman-Gotoh GPU implementation (Liu et al., 2013). It is likely that such optimizations will appreciably improve A21’s throughput.

In an effort to keep up with the increasing amount of sequence data used in clinical and research settings, the usual approach to designing read alignment software has been to focus on increasing throughput. Experience with both CPU-based and GPU-based aligner implementations suggests that the most expeditious way to improve throughput is to add additional computational hardware, that is, to compute read alignments concurrently in multiple threads of execution. In this regard, therefore, GPU hardware is an attractive platform for high-throughput sequence-alignment implementations.

Nonetheless, the highly data-parallel nature of GPU hardware makes it difficult to re-use CPU-based techniques in a GPU implementation. A different approach to exploiting GPU parallelism is to use it for computational tasks that are particularly well suited to the hardware, that are difficult to perform efficiently on sequential CPU threads, and that can improve throughput and accuracy. We see A21 as a first step in this direction.

## ACKNOWLEDGEMENTS

We are grateful to David Luebke and Cliff Wooley of NVidia Corporation for their help in understanding some of the nuances of NVidia GPU hardware.

*Conflict of interest*: none declared.

## REFERENCES

Altschul SF, et al. (1990) Basic Local Alignment Search Tool. J Mol Biol 215, 403–410.

Anderson M et al. (2011) Considerations when evaluating microprocessor platforms. Proceedings of the 3rd USENIX conference on hot topics in parallelism. USENIX Association Berkeley, CA.

Carriero N and Gelernter DH. (1990) How to write parallel programs: a first course. MIT Press, Cambridge, MA. ISBN 9-262-03171-X.

Chen Y, Souaiaia T, Chen, T. (2009) PerM: efficient mapping of short sequencing reads with periodic full sensitive spaced seeds. Bioinformatics 25 (19), 2514–2521.

Genome Reference Consortium. (2014) Human Build 37 patch release 5 (GRCh37.p5). http://www.ncbi.nlm.nih.gov/assembly/GCF_000001405.17/

Glenn TC. (2011) Field guide to next-generation DNA sequencers. Molecular Ecology Resources 11, 759–769.

Gotoh O. (1982) An improved algorithm for matching biological sequences. J Mol Biol 162, 705–708.

Hayden EC. (2014) Is the $1,000 genome for real? Nature News & Comment. http://www.nature.com/news/is-the-1-000-genome-for-real-1.14530, downloaded April 2014.

Holtgrewe M. (2010). Mason – a read simulator for second generation sequencing data. Technical Report TR-B-10-06, Institut für Mathematik und Informatik, Freie Universität Berlin.

Khajeh-Saeed A et al. (2010) Acceleration of the Smith-Waterman algorithm using single and multiple graphics processors. J Computational Physics 229, 4247–4258.

Langmead B and Salzberg S. (2012) Fast gapped-read alignment with Bowtie 2. Nature Methods 9, 357–359.

Li G. et al. (2009) The YH database: the first Asian diploid genome database. Nucleic Acids Research 37(Database issue), D1025–8.

Li H. (2013) Aligning sequence reads, clone sequences and assembly contigs with BWA-MEM. arXiv:1303.3997v1.

Li H et al. (2008) Mapping short DNA sequencing reads and calling variants using mapping quality scores. Genome Research 18, 1851–1858.

Lipman DJ and Pearson WR. (1985) Rapid and sensitive protein similarity searches. Science 227 (4693), 1435–1441.

Liu Y et al. (2013) CUDASW++ 3.0: accelerating Smith-Waterman protein database search by coupling CPU and GPU SIMD instructions. BMC Bioinformatics 14, 117.

Luo R et al. (2013) SOAP3-dp: Fast, Accurate and Sensitive GPU-Based Short Read Aligner. PLoS ONE 8(5): e65632.

Manavski SA and Valle G. (2008) CUDA compatible GPU cards as efficient hardware accelerators for Smith-Waterman sequence alignment. BMC Bioinformatics 9 (Suppl 2), S10.

NVidia Corporation. (2014) NVBIO. http://nvlabs.github.io/nvbio, downloaded May 2014.

SAM/BAM Format Specification Working Group. Sequence Alignment/Map Format Specification (October 18, 2013). https://github.com/samtools/hts-specs, downloaded October 29, 2013.

Schatz MC et al. (2007) High-throughput sequence alignment using Graphics Processing Units. BMC Bioinformatics 8, 474.

Smith TF and Waterman MS. (1981) Identification of common molecular sub-sequences. J Mol Biol 147, 195–197.

Wu TD and Nacu S. (2010) Fast and SNP-tolerant detection of complex variants and splicing in short reads. Bioinformatics 26 (7), 873–881.

